# Acoustic Correlates of the Syllabic Rhythm of Speech: Modulation Spectrum or Local Features of the Temporal Envelope

**DOI:** 10.1101/2022.07.17.500382

**Authors:** Yuran Zhang, Jiajie Zou, Nai Ding

## Abstract

The speech envelope is considered as a major acoustic correlate of the syllable rhythm since the peak frequency in the speech modulation spectrum matches the mean syllable rate. Nevertheless, it has not been quantified whether the peak modulation frequency can track the syllable rate of individual utterances and how much variance of the speech envelope can be explained by the syllable rhythm. Here, we address these problems by analyzing large speech corpora (>1000 hours of recording of multiple languages) using advanced sequence-to-sequence modeling. It is found that, only when averaged over minutes of speech recordings, the peak modulation frequency of speech reliably correlates with the syllable rate of a speaker. In contrast, the phase-locking between speech envelope and syllable onsets is robustly observed within a few seconds of recordings. Based on speaker-independent linear and nonlinear models, the timing of syllable onsets explains about 13% and 46% variance of the speech envelope, respectively. These results demonstrate that local temporal features in the speech envelope precisely encodes the syllable onsets but the modulation spectrum is not always dominated by the syllable rhythm.

## Introduction

In the speech, information is hierarchically organized into units of different sizes, including, e.g., phonemes, syllables, morphemes, words, and many other levels of units such as phrases and sentences. The syllable is a mesoscale speech unit whose time scale lies in between phonemes, the basic phonetic unit, and morphemes, the basic semantic unit. In the last 2 decades, the neural mechanisms underlying the neural representation of syllables have received a significant amount of attention: It is hypothesized that theta-band neural oscillations (4-8 Hz) provide a potential mechanism to parse a continuous speech stream into discrete syllabic units (Assaneo & Poeppel, 2018; Ghitza, 2013; Greenberg, 1999; Hovsepyan, Olasagasti, & Giraud, 2020; Hyafil, Fontolan, Kabdebon, Gutkin, & Giraud, 2015; D. Poeppel, Idsardi, & van Wassenhove, 2008). According to this hypothesis, the syllable is a fundamental unit that transforms an auditory representation of speech into a linguistic representation. This hypothesis is mainly motivated by two observations. First, syllables are perceptually salient units (Liberman, Shankweiler, Fischer, & Carter, 1974; Mehler, Dommergues, Frauenfelder, & Segui, 1981) and are believed to have a reliable acoustic correlate, i.e., the speech envelope (Greenberg, Carvey, Hitchcock, & Chang, 2003). The speech envelope describes the low-frequency fluctuation in sound power (typically below 16 Hz, (Rosen, 1992)), a basic acoustic cue that is well represented in the auditory cortex (Shamma, 2001) and is crucial for speech intelligibility (Taishih Chi, Gao, Guyton, Ru, & Shamma, 1999; T. Chi, Ru, & Shamma, 2005; Elhilali, Chi, & Shamma, 2003; Elliott & Theunissen, 2009). In contrast to the highly variable and context-dependent acoustic features of individual phonemes, the speech envelope provides a simple, articulatory-free cue for syllable boundaries (Stevens, 2002). It has been proposed that the brain may first determine the boundaries between syllables by analyzing the speech envelope, before further decoding the phonetic content (Doelling, Arnal, Ghitza, & Poeppel, 2014).

Second, the mean syllabic rate is between 4 and 8 Hz across languages (Pellegrino, Coupe, & Marsico, 2011), and the peak frequency of the speech modulation spectrum, i.e., the power spectrum of the speech envelope, also lies between 4-8 Hz across languages (Ding et al., 2017; Varnet, Ortiz-Barajas, Erra, Gervain, & Lorenzi, 2017). More importantly, when listening to speech, it has been demonstrated that low-frequency (<∼10 Hz) cortical activity tracks the speech envelope (Ding & Simon, 2012; Lalor & Foxe, 2010; Luo & Poeppel, 2007). Since the speech envelope is considered as a major acoustic correlate of the syllable rhythm, the theta-band envelope-tracking response is often interpreted as a neural marker for the syllabic processing (Doelling et al., 2014; Oganian & Chang, 2019). Since the envelope-tracking response is highly reliable, it has been widely employed to probe, e.g., the neural mechanism to encode acoustically degraded speech (Howard & Poeppel, 2010; Nourski et al., 2009; Zoefel & VanRullen, 2016) and to characterize the speech processing in special populations (Leong & Goswami, 2014).

Although the speech envelope is considered as a major acoustic correlate of the syllabic rhythm, a close look at the literature reveals that this relationship is less well established than typically assumed. For example, the syllable rate varies between 4 and 8 Hz across speakers, speaking styles, and languages (David Poeppel & Assaneo, 2020), but the peak frequency of the modulation spectrum is insensitive to speaking styles and languages (Ding et al., 2017; Varnet et al., 2017). To our knowledge, it has not been tested whether the peak modulation frequency can track the syllable rate, e.g., whether an utterance with a higher syllable rate shows a higher peak modulation frequency. Furthermore, the modulation spectrum of speech peaks around 4-5 Hz only when a 1/f trend is removed. Otherwise, the peak modulation frequency is around 0 Hz. In contrast, no 1/f trend needs to be removed when calculating the syllable rhythm. Even more importantly, sharing the same center frequency does not guarantee that two rhythms are synchronized.

The current study quantifies the relationship between syllables and speech envelope based on large speech corpora (>1000 hours of recording). Furthermore, on top of the traditional modulation spectrum analysis, we employed a linear temporal response function (TRF) and a more advanced deep neural network (DNN) to model the relationship between speech envelope and syllable onsets. In the following, we first quantified whether the peak modulation frequency was correlated with the mean syllable rhythm across individual speakers. In this analysis, we considered several methods to calculate the peak modulation frequency, e.g., based on the broadband or narrowband modulation spectrum, and several methods to calculate the syllable rhythm, e.g., the mode of syllable distribution, the syllable rate, or the articulation rate. Next, we analyzed the time locking between speech envelope and syllable onsets using the TRF, which could extract the time-domain envelope features associated with a syllable onset and also reveal how much percent of the variance of speech envelope can be explained by syllable onsets. The TRF is a simple and interpretable model but it can only characterize Linear Time-Invariant (LTI) relationships between two signals. Therefore, we also employed an advanced DNN model to better characterize the more complex relationship between the speech envelope and syllable onsets.

## Results

### Relationship between modulation spectrum and syllable rhythm

We first compared the shape of the speech modulation spectrum and the distribution of syllable duration. We separately extracted the broadband and narrowband speech envelope (Figs 1A and 1B) and transformed them into the frequency domain to obtain the broadband and narrowband modulation spectrum (Fig 1C). The modulation spectrum was averaged with each corpus and showed a peak around 4 Hz for both broadband and narrowband envelope. The syllable rhythm was characterized using 3 measures (Fig 2A). The first measure was the mode of the histogram of the reciprocal of the duration of individual syllables (Greenberg et al., 2003). The second measure was the syllable rate, i.e., the number of syllables divided by the duration of the recording, and the third measure was the articulation rate, i.e., the number of syllables divided by the total duration of syllables. The difference between the syllable rate and articulation rate was that the former measure considered the silence periods in speech but the latter measure did not. All three measures of syllable rhythm coincide with the peak frequency of the modulation spectrum. The differences between peak modulation frequency and the syllable rhythm measures are shown in Fig 3B.

**Fig 1.**
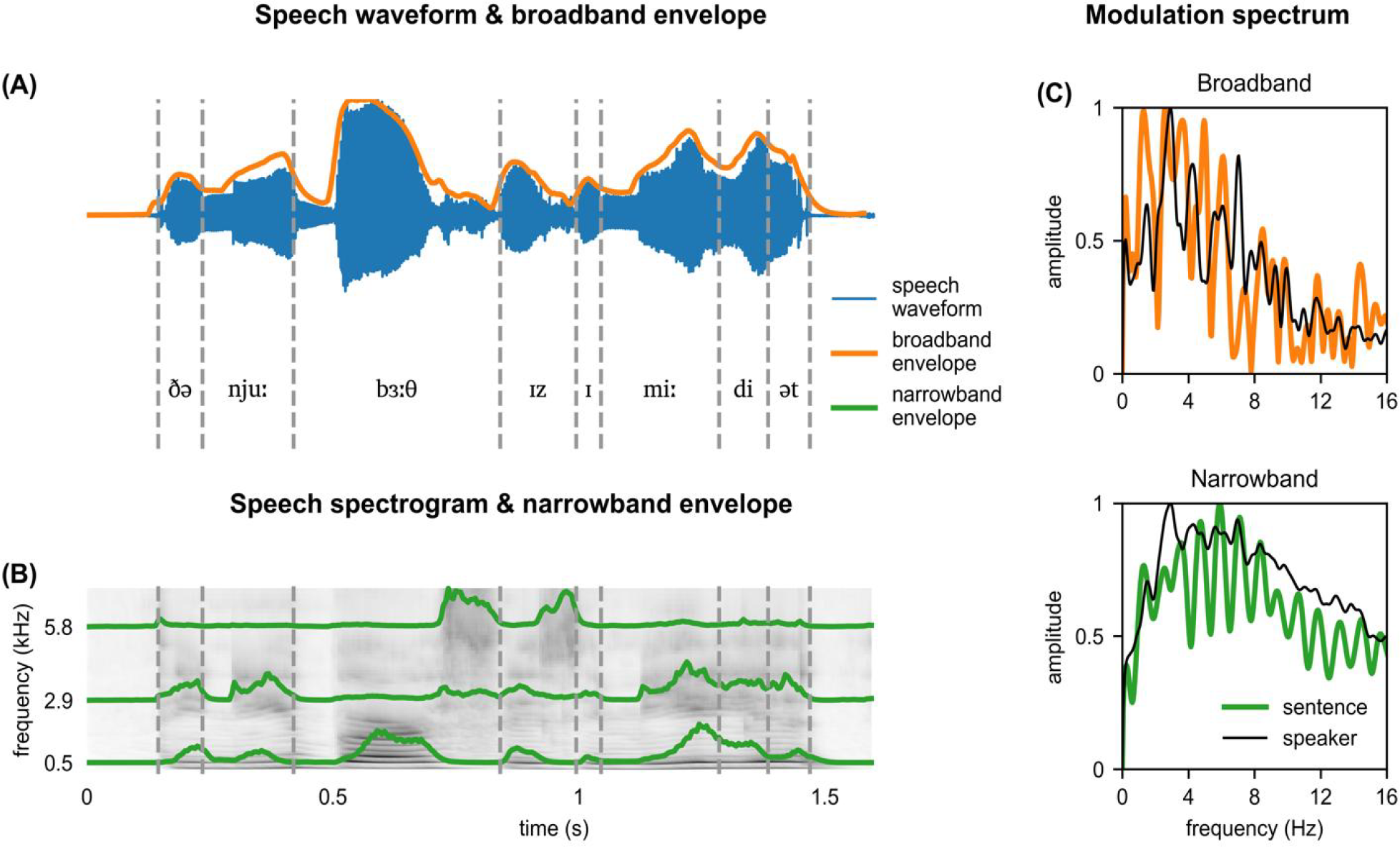
Speech envelope and modulation spectrum. (A) The speech waveform and broadband envelope of a sentence (“the new birth is immediate”) from a speaker. Syllable boundaries are marked by dashed lines. (B) Speech spectrogram and narrowband envelope. Each row in the spectrogram constitutes a narrowband envelope. (C) Normalized modulation spectrum. The broadband and narrowband modulation spectra are calculated based on the broadband and narrowband envelope, respectively. The colored and black lines are calculated based on that sentence and all sentences of that speaker, respectively.

**Fig 2.**
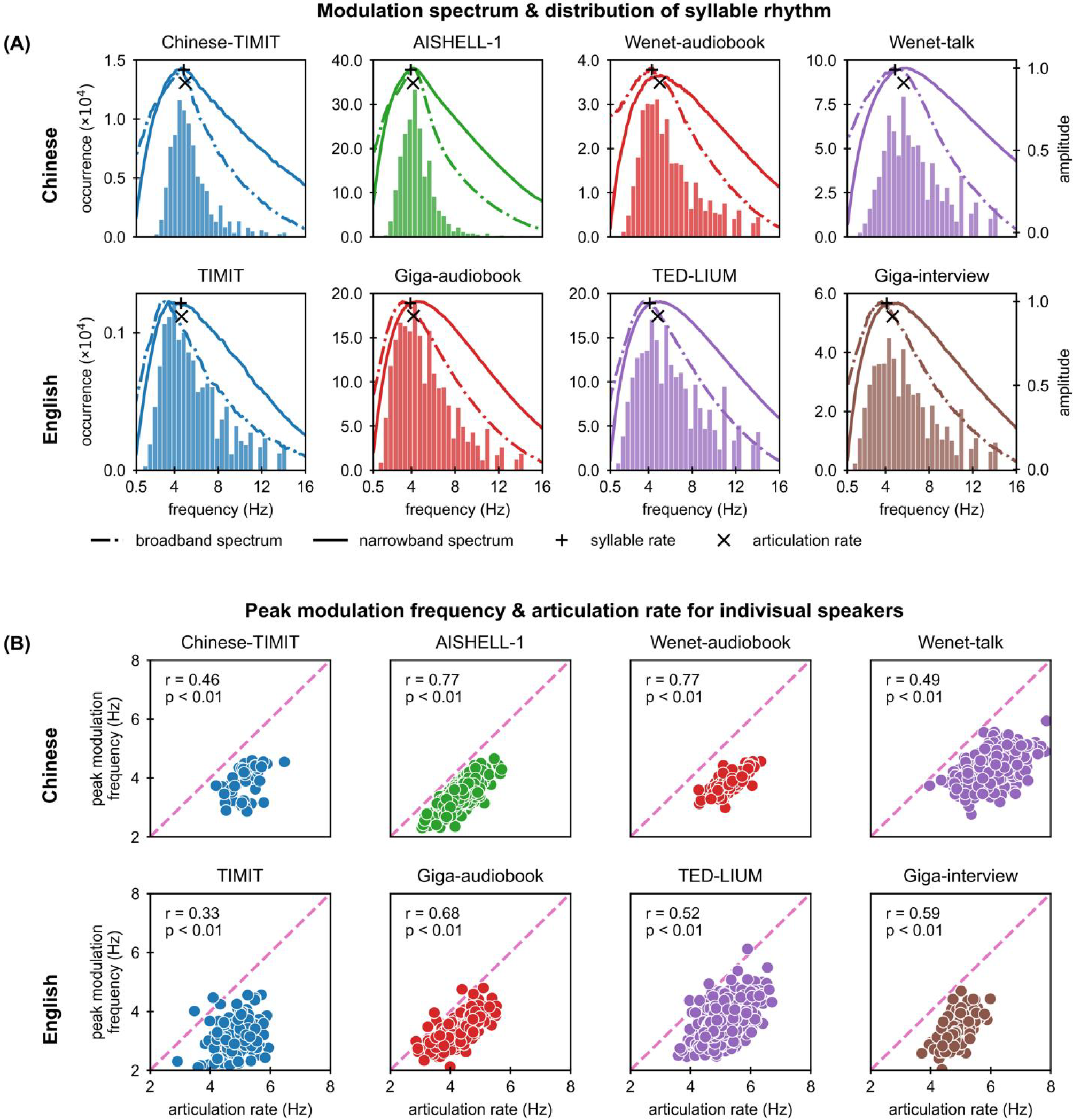
Peak modulation frequency, syllable rhythm, and their correlation. (A) The broadband and narrowband modulation spectra, as well as the histogram of the reciprocal of syllable duration, are shown for each speech corpora. The mean syllable rate and the mean articulation rate are shown by markers. For each spoken language, each corpus is shown with a unique color that is consistent in all figures. (B) Correlation between peak modulation frequency and articulation rate for individual speakers. The peak modulation frequency is extracted from the broadband modulation spectrum using Method 1.

**Fig 3.**
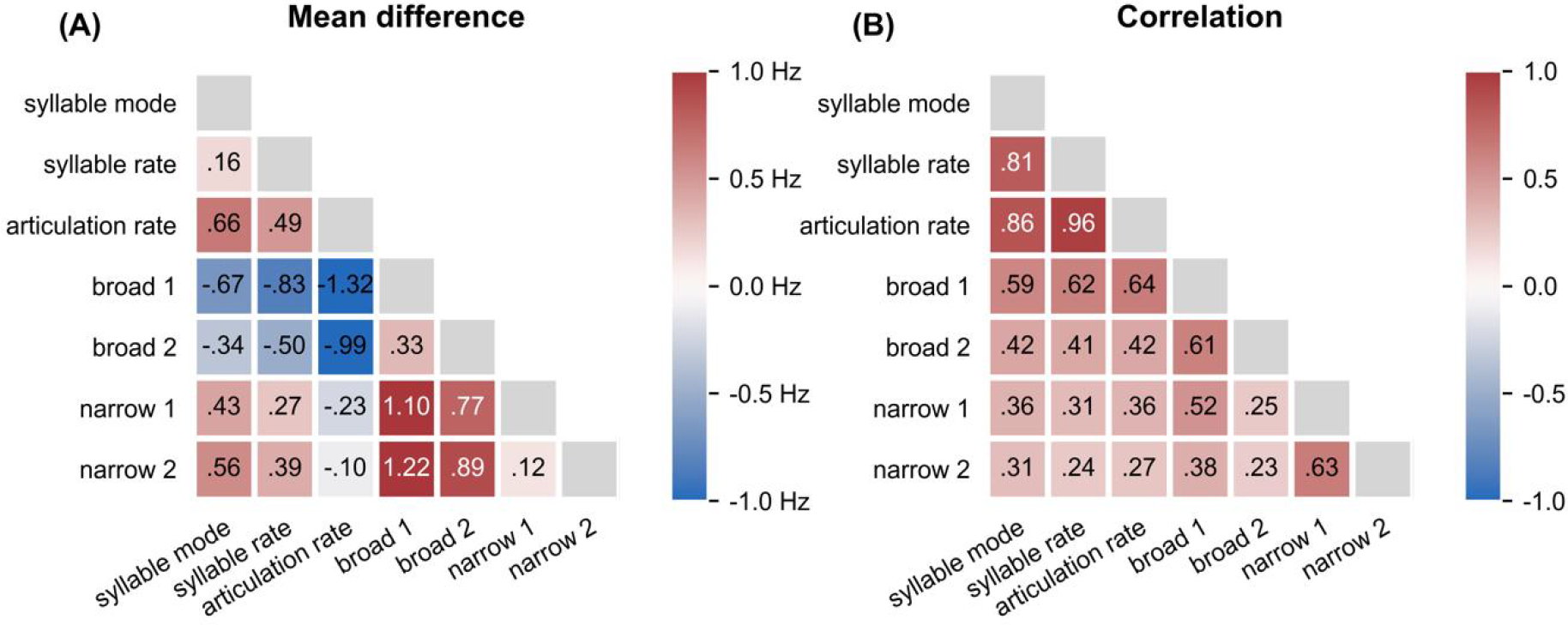
Mean difference and correlation between different measures describing the syllable rhythm and peak modulation frequency. (A) Each measure is averaged across corpora and the difference between measures is shown. (B) Pairwise correlation of different methods is calculated for each corpus, and the correlation coefficient of each corpus is weighted by the total duration of the corpus and averaged.

Next, we asked whether the syllable rhythm and peak modulation frequency were correlated across individual speakers. Since each corpus had multiple sentences from a speaker and we applied two methods to extract the mean peak frequency across the sentences (see Methods). Both the broadband and narrowband modulation spectra were considered in this analysis. The peak modulation frequency calculated using either method correlated with the syllable mode, as well as the syllable rate and the articulation rate of the speaker. The correlation coefficients between different measures of speech modulation and syllable rhythm were averaged across corpora and were shown in Fig 3A. The peak frequency of the broadband modulation spectrum calculated using Method 1 was most correlated with the syllable rhythm. Therefore, in the following, we restricted our discussion to the broadband modulation spectrum and employed Method 1 to integrate the peak modulation frequency across sentences. Since the three measures of syllable rhythm similarly correlated with the peak modulation frequency, in the following, we only considered the articulation rate, i.e., 1 over the mean syllable duration.

We further illustrated the relationship between peak modulation frequency and the articulation rate in Fig 2B, for individual corpora. The correlation was the lowest for the two smaller corpora, i.e., TIMIT and Chinese-TIMIT, and the highest for AISHELL. Chinese-TIMIT and AISHELL both contained read Chinese sentences, and mainly differed in the recording duration per speaker. To test whether recording duration influenced the correlation between peak modulation frequency and articulation rate, we calculate the correlation based on subsets of the recordings. It was observed that the correlation increased when the recording duration of individual speakers increased (Fig 4). When the recording duration per speaker was 4 s, the correlation between peak modulation frequency and articulation rate was only around 0.25 while the asymptote correlation was above 0.5 for most corpora. Based on a sigmoid fit, the correlation reached 95% of its maximum when the recording duration was beyond 512 seconds for most corpora.

**Fig 4.**
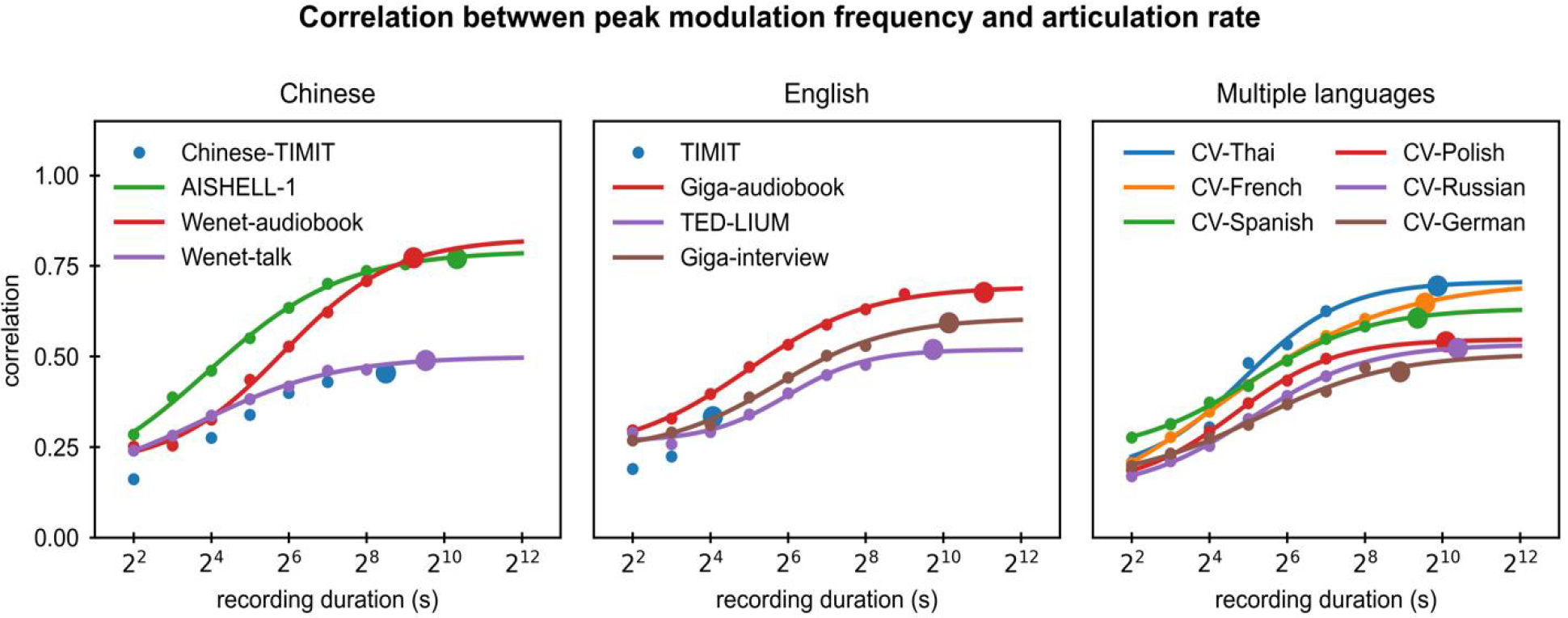
Correlation between peak modulation frequency and articulation rate as a function of recording duration. The recording duration is the duration of the speech signal available for each speaker. The smaller and bigger markers separately show the correlation calculated based on a subset of the speech corpus or the whole corpus. The result from each large corpus is a fit by the sigmoid function.

### Temporal mapping between speech envelope and syllable onsets

The correlation between peak modulation frequency and the syllable rhythm characterized whether the envelope rhythm and the syllable rhythm fell into the same frequency range. Having the same frequency range, however, did not indicate that two sequences were time-locked. Next, we used the temporal response function (TRF) to analyze the degree of phase-locking between speech envelope and syllable onsets. The syllable onsets were represented by a 0-1 sequence whose value was 1 only at the onset of a syllable (Fig 5A). The broadband envelope was modeled by convolving the syllable onset sequence with a TRF. For the two smaller corpora, i.e., TIMIT and Chinese-TIMIT, the TRF was estimated based on the whole corpora, yielding a speaker-independent model. For other larger corpora, the TRF was separately estimated for each speaker, yielding a speaker-dependent model.

**Fig 5.**
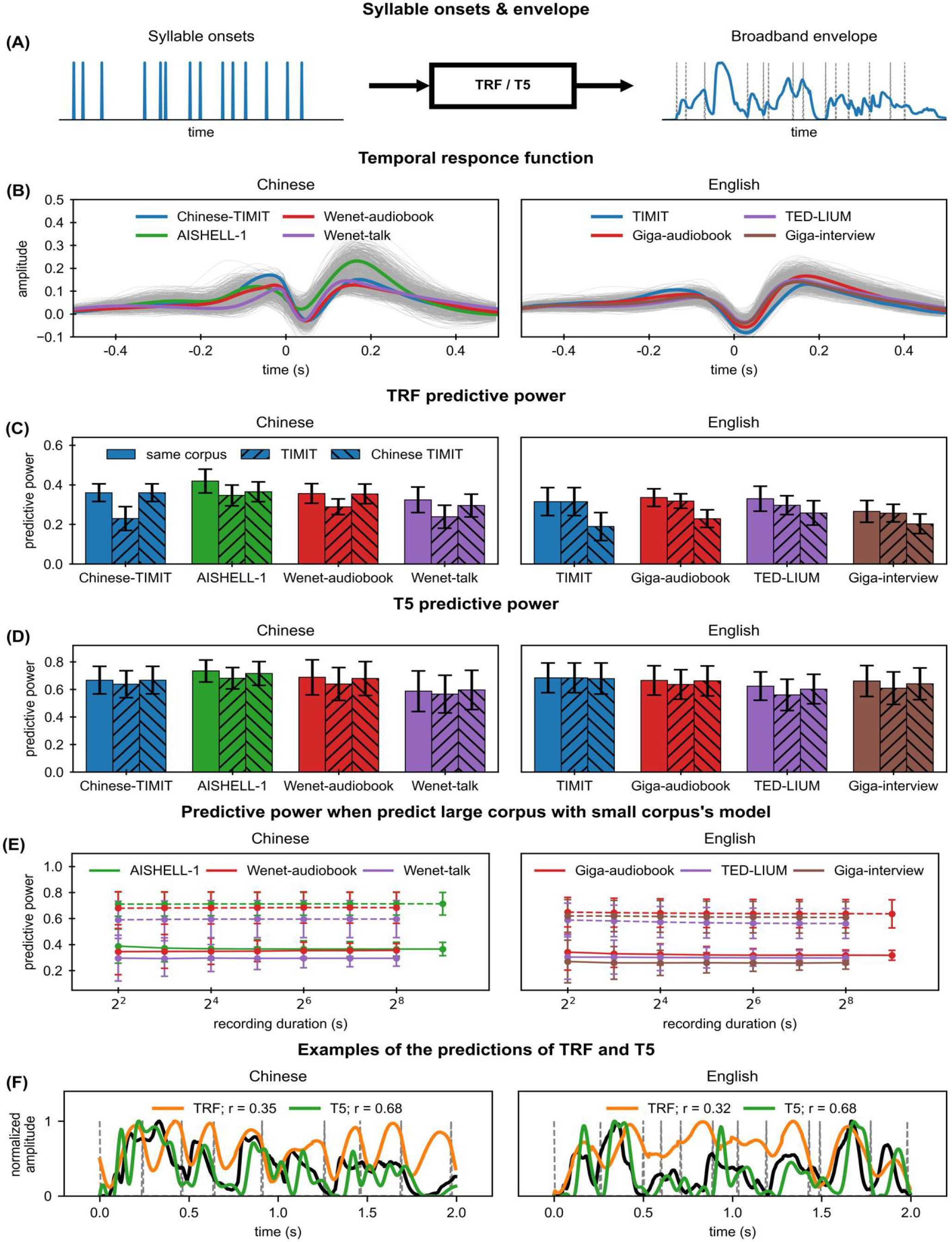
TRF and predictive power. (A) syllable onset sequence and the broadband envelope, which are the input and output of the TRF or T5 model, respectively. (B) TRF for each corpus. The mean TRF averaged over each corpus is color-coded. For each large corpus, the TRF for each speaker is shown by a gray curve. (C) The predictive power of TRF. Each bar shows one standard deviation. The three bars for each corpus show results when the model is trained on different corpora (see legend). (D) The predictive power of T5. (E) The predictive power of TRF (solid lines) or T5 (dashed lines) when predicting a large corpus with a small corpus’s model as a function of recording duration. (F) The examples of the predictions of the TRF and T5, which are cut off at 2 s. The sentences are randomly picked from Chinese-TIMIT and TIMIT with the constraint that the predictive power of the sentence does not strongly deviate from the mean predictive power of the corpus for both models. Syllable onsets and broadband envelopes are marked by the gray dashed lines and black solid lines, respectively.

The predictive power of TRF, i.e., the correlation coefficient between the actual envelope and the envelope predicted by syllable onsets, mainly ranged between 0.3 and 0.5 for individual speakers (Fig 5C). The square of the predictive power for each corpus was 12.56 ± 5.23%, which could be interpreted as the percent of envelope variance explained by the syllable onset. The TRF showed a trough at 32 ± 8 ms latency after a syllable onset (Fig 5B), indicating a local minimum in the broadband envelope after a syllable onset. To analyze whether a long recording was required to reveal the phase-locking between speech envelope and syllable onsets, we calculated the predictive power of the speaker-independent TRFs on subsets of large corpora. It was observed that the TRF learned based on the two smaller corpora could generalize well to the larger corpora, and the predictive power was above chance (Fig 5C). More importantly, the predictive power did not clearly change with the recording duration (Fig 5D). Furthermore, the TRF estimated based on Chinese could predict the relationship between syllable onsets and envelope above chance for English, and vice versa (Fig 5C). The results suggested that TRF predictive power was more robust to recording duration and could generalize to more testing data even across languages.

The TRF is a LTI system model. Nevertheless, the mapping between syllable onsets and speech envelope is certainly time-variant: The envelope of each syllable is not the same. Therefore, we also employed a DNN model to better estimate the upper limit of how well syllable onsets can explain the speech envelope. The DNN model we employed is T5 (Raffel et al., 2020), a state-of-the-art model for sequence-to-sequence modeling. The predictive power of the T5 ranged from roughly 0.6 to 0.8, almost twice the predictive power of the TRF (Fig 5C). Furthermore, Similar to the TRF model, the T5 model generalizes well across languages and is robust to the duration of the speech recording being analyzed.

## Discussion

The current study investigates the relationship between speech envelope and syllable boundaries. In contrast to the common belief that the narrowband speech envelope is a main carrier of the syllable rhythm, the study shows that the peak frequency of the broadband envelope is more strongly correlated with the syllable rate, suggesting that the syllable rhythm is better reflected by coherent fluctuations across frequencies. Furthermore, the broadband peak modulation frequency is only clearly correlated with the syllable rate when averaged over several minutes of speech recordings. In contrast, the TRF/T5 analyses reveal that a component of the broadband speech envelope is reliably phase-locked to the onset of individual syllables.

The TRF model assumes that all syllables have the same broadband speech envelope, i.e., the same duration, amplitude, and time course of the envelope, regardless of the phonetic content and the context. Although this assumption is clearly oversimplistic, based on the TRF model, the onset of individual syllables explains about 13% of the variance of the broadband speech envelope. Furthermore, when considering contextual information, i.e., when the envelope of a syllable is predicted based on the onset time of all syllables in a sentence instead of just the onset time of the current syllable, syllable onsets can explain about 46% of the variance of the speech envelope based on a DNN model, i.e., T5. This ratio is quite high in the sense that the models do not consider the phonetic content of speech, which can certainly influence the shape of the speech envelope. For example, the syllables ‘oh’ and ‘skirt’ certainly have distinct speech envelopes, even when they have equal duration. Finally, both the TRF and DNN models generalize well between English and Chinese, two languages that have distinct phonetic properties.

These results suggested that the narrowband modulation spectrum potentially reflects intrinsic dynamic properties of the articulators, while the broadband envelope is more strongly influenced by the timing of syllable onsets. Even for the broadband envelope, the overall periodicity, i.e., the peak modulation frequency, is not reliably correlated with the syllable rate within relatively short time intervals that correspond to the duration of a typical sentence, possibly attributable to the influence of phonetic content and word/phrase-level prosodic information. These results are not surprising in the sense that the speech envelope is known to possess strong power at very low modulation frequencies well below the syllable rate. The general 1/f trend in the modulation spectrum can render the syllable-rate peak frequency highly variable across utterances, even after the 1/f trend is corrected. In summary, the local time course of the broadband envelope provides a reliable cue for syllable onsets, as is revealed by the TRF analysis, but the mean periodicity of the broadband envelope provides a reliable cue for the mean syllable rate only when averaged over minutes of recordings.

### Speech Modulation Spectrum

The narrowband modulation spectrum has been widely applied to characterize the rhythm of speech. Although the narrowband spectrum is similar in shape across languages, speakers, and speaking styles (Ding et al., 2017; Varnet et al., 2017), it still could be affected by language (Varnet et al., 2017) and speaking style (Bosker & Cooke, 2018; Krause & Braida, 2004). Summarizing the results in previous studies, the mean peak frequency is 4.5 Hz for the English telephone corpus Switchboard (Greenberg et al., 2003), between 4.4 and 4.8 Hz for English corpora with different speaking styles (Ding et al., 2017), between 4.3 and 5.4 Hz for naturalistic discourse-level speech across nine languages (Ding et al., 2017), and between 4.3 and 5.5 Hz for semi-read speech (SRS) corpus in ten languages (Varnet et al., 2017). In the current study, the peak frequency of the narrowband modulation spectrum is between 4.2 and 5.9 Hz across corpora and was on average 4.9 Hz. The broadband modulation spectrum also shows a peak, and the peak frequency ranges from 3.5 to 4.5 Hz across six languages (David Poeppel & Assaneo, 2020). In the current study, the peak frequency of the broadband modulation spectrum is between 3.3 and 5.0 Hz across corpora and was on average 4.3 Hz. The peak frequency of the broadband modulation spectrum is lower than that for the narrowband modulation spectrum in both the current study and previous studies (David Poeppel & Assaneo, 2020). Therefore, for both the broadband and narrowband modulation spectrum, the peak frequency is highly consistent across languages and speaking styles and the variation across corpora is within 1.7 Hz.

### Syllable Rhythm

In contrast to the highly consistent peak frequency for the modulation spectrum, the syllable rate reported in the literature greatly varies across languages. The mean syllable rate is between 5.2 and 7.8 Hz across eight languages in one study (Pellegrino et al., 2011) and is between 4.3 and 9.1 Hz across seventeen languages in another study (Coupé, Oh, Dediu, & Pellegrino, 2019). In the current study, the syllable rate is between 3.5 and 6.1 Hz across eight languages. Similarly, based on the literature, the articulation rate greatly varies across speaking styles. It is between 3.1 and 5.6 Hz for different speaking styles in English (Jacewicz, Fox, O’Neill, & Salmons, 2009) and is between 3.8 and 7.2 Hz in German (Jessen, 2007). In the current study, the articulation rate is between 4.3 and 5.2 Hz across speaking styles for English and between 4.3 and 6.0 Hz for Chinese. Finally, the syllable mode is only previously reported for Switchboard, i.e., 5.2 Hz (Greenberg et al., 2003), and varies between 3.5 and 5.3 Hz across corpora in the current study. In summary, for different languages, the syllable rate varies between 3.5 and 9.1 Hz. For speaking styles, the articulation rate varies between 3.1 and 7.2 Hz across corpora.

The results reported in the literature are based on different corpora, which makes it challenging to compare the results across studies. For example, the articulation rate is by definition faster than the syllable rate, while the articulation rate reported in the literature is often slower than the syllable rate. A likely reason is that the studies reporting high syllable rates are studies that ask the speaker to read materials after familiarizing (Coupé et al., 2019). The comparison between languages is especially challenging given the strong influence of speaking style. For example, when comparing the syllable rhythm in Chinese and English, a study that analyzed utterances that are read clearly or naturally reveals a faster articulation rate for Chinese (Ann Burchfield & Bradlow, 2014), while studies that analyzed utterances of familiar sentences reveal a faster syllable rate for English (Coupé et al., 2019; Pellegrino et al., 2011). The current study analyzed two corpora for reading Chinese sentences and one corpus for reading English sentences, while the syllable rhythm of the English corpus is in between of the syllable rhythms of the two Chinese corpora.

Finally, the syllable rhythms reported in the literature show much greater variance than the peak modulation frequencies. Similarly, in the current study, when the two measures are calculated based on the same set of corpora, the variance across corpora tends to be larger for the syllable rhythm. However, the variance across speakers in each corpus tends to be smaller for the syllable rate than the peak modulation frequencies.

### Relationship between Modulation Spectrum and Syllable Rhythm

Although it was commonly assumed that the modulation spectrum reflected the distribution of syllable rate (Greenberg, Hollenback, & Ellis, 1996), here it was demonstrated that the peak frequency of the narrowband modulation spectrum only weakly correlated with the syllable rate of a speaker (correlation coefficient below 0.4). For the broadband modulation spectrum, the correlation coefficient can reach 0.6, but observing such a correlation requires a very long speech recording per speaker, e.g., 512 s. The higher correlation of the broadband modulation spectrum suggests that the syllable rate is more strongly related to the acoustic fluctuations that are coherent across frequency bands, compared to the acoustic fluctuations in individual frequency bands. When only 4 s of speech recording is available for each speaker, which corresponds to the duration of a relatively long sentence, the correlation between broadband peak modulation frequency and syllable rate is also weak (correlation coefficient around 0.25). Therefore, the peak modulation frequency of speech does not precisely capture the syllable rate of an individual speaker. The peak modulation frequency is typically lower than the syllable rate, possibly attributable to the influence of phrasal level prosodic information.

The reliability of the speech modulation frequency, however, does not indicate that it simply lacks sensitivity to sound rhythms. For example, the peak frequency is consistently different between speech and western music. Furthermore, if a speech signal is artificially time-compressed, the speech envelope is time-compressed accordingly and the modulation frequency shifts (S3 Fig). Therefore, the current results suggest that utterances of different syllable rates are not simply time-scaled versions of each other. Instead, these utterances still share some fundamental acoustic rhythm as characterized by the narrowband modulation spectrum, which possibly originates from the biophysical properties of the human articulator (Chandrasekaran, Trubanova, Stillittano, Caplier, & Ghazanfar, 2009). Previous studies have also widely documented that an increase in the speech rate is accompanied by, e.g., vowel reduction (Lindblom, 1963; Taylor, Theobald, & Matthews, 2014), consonant reduction (van Son & Pols, 1999), an increase in co-articulation (Agwuele, Sussman, & Lindblom, 2008), and lightly changes in the consonant/vowel ratio (Kessinger & Blumstein, 1998).

### Relationship between Speech Envelope and Syllable Onsets

The TRF analysis reveals that, even with a short 4-s speech recording, the broadband envelope of speech shows reliable phase locking to the onset of individual syllables. The TRF shows that the envelope tends to show a local minimum at about 32 ms after a syllable onset and peaks at about 160 ms after a syllable onset. The results are consistent with previous findings that the syllable boundaries can be extracted from the speech envelope (Prasanna, Reddy, & Krishnamoorthy, 2009; D. Wang & Narayanan, 2007), and that a syllable typically has one local maximum in the envelope, which corresponds to the nucleus (Greenberg et al., 2003; Hooper & Bybee, 1976), while the boundaries of syllables more closely correspond to troughs in the envelope (Mermelstein, 1975). The current study, however, extends the previous studies by quantifying how much percent of the variance of the speech envelope can be explained by syllable onsets. Furthermore, previous studies that employ the speech envelope to extract syllable boundaries often view the results as correct if the extracted boundary is within 40 ms (Kamper, Jansen, & Goldwater, 2016) or 50 ms (Villing, Ward, & Timoney, 2006) from the human-annotated boundaries, while the current study quantifies the timing between the troughs in envelopes and the syllable onsets.

## Materials and Methods

### Speech materials

Seven speech corpora were included in analysis, i.e., DARPA-TIMIT (Garofolo, Lamel, Fisher, Fiscus, & Pallett, 1993), Chinese-TIMIT (Yuan, Ding, Liao, Zhan, & Liberman, 2017), Aishell-1 (Bu, Du, Na, Wu, & Zheng, 2017), WenetSpeech (Zhang et al., 2021), GigaSpeech (Chen et al., 2021), TED-LIUM (Rousseau, Deléglise, & Esteve, 2012), and Common-voice (Ardila et al., 2020). All corpora had transcriptions available. One corpus, i.e., Common-voice, had multiple languages and we selected six languages that the mean speaker duration is longer than 512s, i.e., Thai, French, Spanish, Polish, Russian, and German. For corpora containing discourse-level recordings, we separated the recordings into sentences based on the sentence boundaries provided in the corpora. All sentences were resampled to 16 kHz and converted into a single channel. All sentences were within 20 s. Corpus processing scripts are available at https://github.com/austin-365/ms-tools.

Two corpora were multi-domain corpora containing multiple speaking styles, i.e., WenetSpeech and GigaSpeech, and for these 2 corpora, we selected three speaking styles for analysis, i.e., audiobook, talk, and interview. The talk category included lectures, e.g., TED talks for English and Yixi talks for Chinese, each talk was given by a single speaker, and the interview category contained recordings from both the interviewer and the interviewees. Audiobook was a category available in both corpora, and talk was a category for WenetSpeech. Audios in the interview category were manually selected from the “youtube” category in GigaSpeech, with the criterion that no background music was present. See Table 2 for details.

**Table 1.**
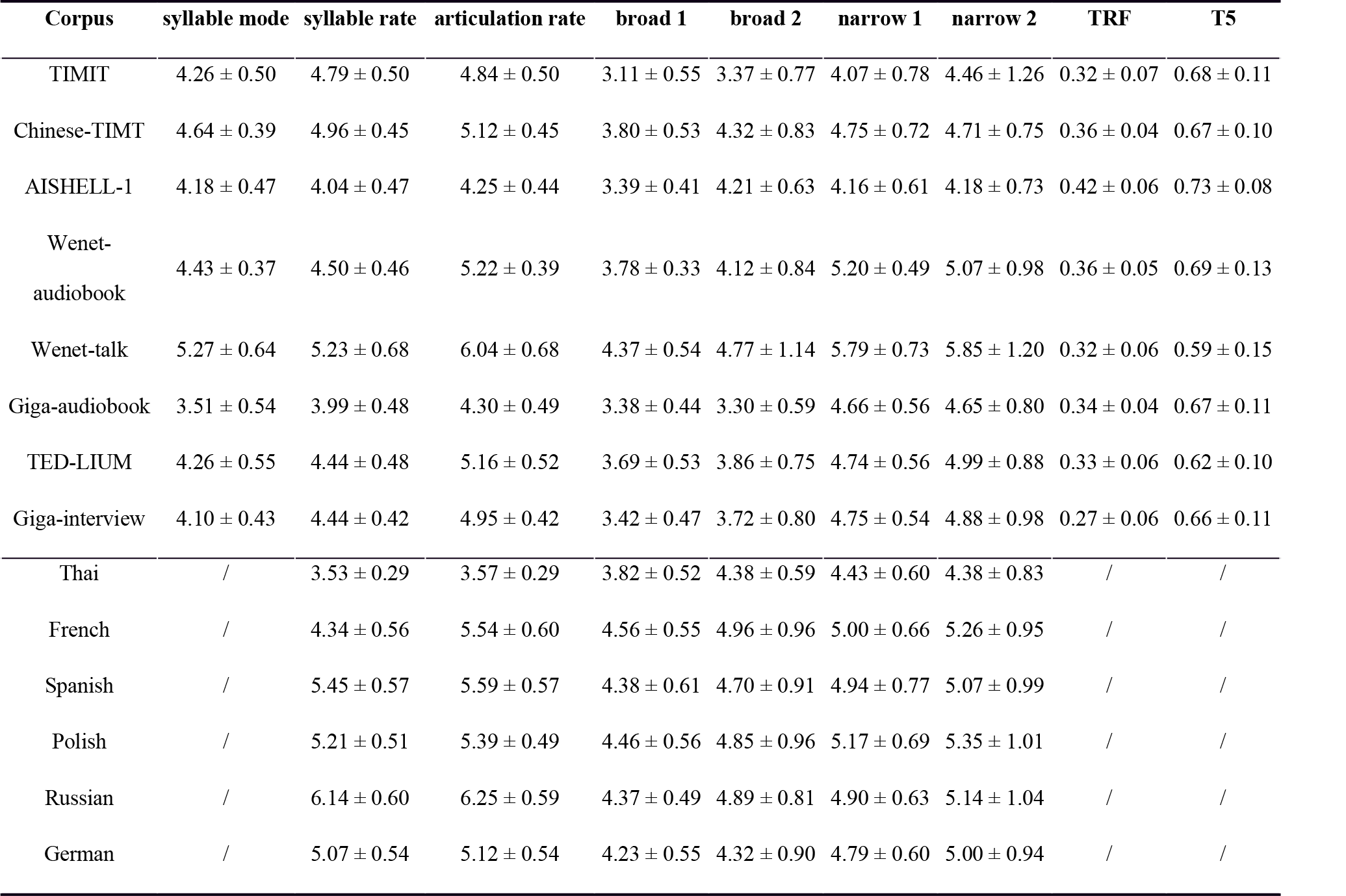
Mean and standard deviation of measures describing the syllable rhythm or modulation rate.

| Corpus | syllable mode | syllable rate | articulation rate | broad 1 | broad 2 | narrow 1 | narrow 2 | TRF | T5 |
| --- | --- | --- | --- | --- | --- | --- | --- | --- | --- |
| TIMIT | 4.26 ± 0.50 | 4.79 ± 0.50 | 4.84 ± 0.50 | 3.11 ± 0.55 | 3.37 ± 0.77 | 4.07 ± 0.78 | 4.46 ± 1.26 | 0.32 ± 0.07 | 0.68 ± 0.11 |
| Chinese-TIMIT | 4.64 ± 0.39 | 4.96 ± 0.45 | 5.12 ± 0.45 | 3.80 ± 0.53 | 4.32 ± 0.83 | 4.75 ± 0.72 | 4.71 ± 0.75 | 0.36 ± 0.04 | 0.67 ± 0.10 |
| AISHELL-1 | 4.18 ± 0.47 | 4.04 ± 0.47 | 4.25 ± 0.44 | 3.39 ± 0.41 | 4.21 ± 0.63 | 4.16 ± 0.61 | 4.18 ± 0.73 | 0.42 ± 0.06 | 0.73 ± 0.08 |
| Wenet-audiobook | 4.43 ± 0.37 | 4.50 ± 0.46 | 5.22 ± 0.39 | 3.78 ± 0.33 | 4.12 ± 0.84 | 5.20 ± 0.49 | 5.07 ± 0.98 | 0.36 ± 0.05 | 0.69 ± 0.13 |
| Wenet-talk | 5.27 ± 0.64 | 5.23 ± 0.68 | 6.04 ± 0.68 | 4.37 ± 0.54 | 4.77 ± 1.14 | 5.79 ± 0.73 | 5.85 ± 1.20 | 0.32 ± 0.06 | 0.59 ± 0.15 |
| Giga-audiobook | 3.51 ± 0.54 | 3.99 ± 0.48 | 4.30 ± 0.49 | 3.38 ± 0.44 | 3.30 ± 0.59 | 4.66 ± 0.56 | 4.65 ± 0.80 | 0.34 ± 0.04 | 0.67 ± 0.11 |
| TED-LIUM | 4.26 ± 0.55 | 4.44 ± 0.48 | 5.16 ± 0.52 | 3.69 ± 0.53 | 3.86 ± 0.75 | 4.74 ± 0.56 | 4.99 ± 0.88 | 0.33 ± 0.06 | 0.62 ± 0.10 |
| Giga-interview | 4.10 ± 0.43 | 4.44 ± 0.42 | 4.95 ± 0.42 | 3.42 ± 0.47 | 3.72 ± 0.80 | 4.75 ± 0.54 | 4.88 ± 0.98 | 0.27 ± 0.06 | 0.66 ± 0.11 |
| Thai | / | 3.53 ± 0.29 | 3.57 ± 0.29 | 3.82 ± 0.52 | 4.38 ± 0.59 | 4.43 ± 0.60 | 4.38 ± 0.83 | / | / |
| French | / | 4.34 ± 0.56 | 5.54 ± 0.60 | 4.56 ± 0.55 | 4.96 ± 0.96 | 5.00 ± 0.66 | 5.26 ± 0.95 | / | / |
| Spanish | / | 5.45 ± 0.57 | 5.59 ± 0.57 | 4.38 ± 0.61 | 4.70 ± 0.91 | 4.94 ± 0.77 | 5.07 ± 0.99 | / | / |
| Polish | / | 5.21 ± 0.51 | 5.39 ± 0.49 | 4.46 ± 0.56 | 4.85 ± 0.96 | 5.17 ± 0.69 | 5.35 ± 1.01 | / | / |
| Russian | / | 6.14 ± 0.60 | 6.25 ± 0.59 | 4.37 ± 0.49 | 4.89 ± 0.81 | 4.90 ± 0.63 | 5.14 ± 1.04 | / | / |
| German | / | 5.07 ± 0.54 | 5.12 ± 0.54 | 4.23 ± 0.55 | 4.32 ± 0.90 | 4.79 ± 0.60 | 5.00 ± 0.94 | / | / |

**Table 2.** speech corpora.

| Name | Source | Language | Speaking Style | # speaker | Speaker Duration (s, mean ± S.D.) | Total Duration (h) |
| --- | --- | --- | --- | --- | --- | --- |
| TIMIT | DARPA-TIMIT | English | Read sentences | 178 | $19.1 \pm 2.5$ | 0.9 |
| Chinese-TIMIT | Chinese-TIMIT | Chinese | Read sentences | 50 | $415.1 \pm 32.9$ | 5.8 |
| AISHELL-1 | AISHELL-1 | Chinese | Read sentences | 400 | $1610.6 \pm 181.2$ | 179 |
| Wenet-audiobook | WenetSpeech | Chinese | Read audiobooks | 135 | $636.6 \pm 201.5$ | 23.9 |
| Wenet-talk | WenetSpeech | Chinese | Talk | 238 | $820.7 \pm 254.9$ | 54.3 |
| Giga-audiobook | GigaSpeech | English | Read audiobooks | 264 | $2303.8 \pm 931.6$ | 168.9 |
| TED-LIUM | TED-LIUM | English | Talk | 646 | $963.6 \pm 157.7$ | 172.9 |
| Giga-interview | GigaSpeech | English | Interviews | 121 | $1214.6 \pm 736.2$ | 40.8 |
| CV-Thai | Common-voice | Thai | Read sentences | 38 | $1434.5 \pm 3024.1$ | 15.1 |
| CV-French | Common-voice | French | Read sentences | 307 | $1188.8 \pm 442.2$ | 101.4 |
| CV-Spanish | Common-voice | Spanish | Read sentences | 174 | $961.1 \pm 466.1$ | 46.5 |
| CV-Polish | Common-voice | Polish | Read sentences | 190 | $1593.2 \pm 3100.6$ | 84.1 |
| CV-Russian | Common-voice | Russian | Read sentences | 226 | $1809.9 \pm 4012.6$ | 113.6 |
| CV-German | Common-voice | German | Read sentences | 400 | $693.9 \pm 146.0$ | 77.1 |

### Envelope and modulation spectrum

The speech envelope and modulation spectrum were extracted using the same methods in (Ding et al., 2017), which were briefly outlined here. The envelope was extracted by using an auditory model that filtered the sound signal into 128 frequency bands. In each band, the filtered sound signal was half-wave rectified, smoothed by a single-pole low-pass filter with a time constant of 8 ms, and decimated to 200 Hz. The output of the auditory model in each frequency band was referred to as the narrowband envelope (*N* = 128), and the average of all 128 narrowband envelopes was referred to as the broadband envelope. The modulation spectrum was obtained by applying the Discrete Fourier Transform (DFT) to the square of the speech envelope. The narrowband spectrum was the root mean square value (RMS) of the 128 modulation spectra corresponding to the 128 narrowband envelopes, and the broadband spectrum was the modulation spectrum of the broadband envelope. The spectrum in Fig 2a was normalized by its maximal value between 0.5 and 16 Hz. In the modulation spectrum analysis, the envelope of each sentence was zero-padded to 20 s so that the frequency resolution for each sentence was the same.

The peak modulation frequency of a speaker was calculated using two methods, based on the modulation spectrum of individual sentences. Method 1 calculated the peak modulation frequency based on each sentence and then average the peak modulation frequency across sentences. Method 2 first calculated the RMS of the modulation spectra across all sentences and then extracted the peak of the RMS spectrum.

### Syllable

The boundary between syllables was automatically extracted and validated based on the corpora for which phoneme boundaries were manually labeled (see Appendix). The boundaries between phonemes were extracted based on the audio and transcription using the Montreal Forced Aligner (MFA, (McAuliffe, Socolof, Mihuc, Wagner, & Sonderegger, 2017)). The MFA also provided the boundaries between words for English and characters for Chinese. For Chinese, the syllable boundary was the same as the boundary between characters, which were available from MFA. For English, the syllable boundary was determined by grouping phonemes into syllables based on a dictionary (Unisyn Lexicon, (Fitt, 2001)). For other languages, the syllable boundary was not extracted, attributable to the diversity in syllable structure. The distribution, i.e., probability density function, of the reciprocal of the syllable duration was estimated by the Gaussian kernel density method with the window width determined using Scott’s rule (Scott, 2015).

### TRF modeling

The phase-locking between speech envelope and syllable onsets was characterized using the Temporal Response Function (Crosse, Di Liberto, Bednar, & Lalor, 2016; Y. Wang, Zhang, Zou, Luo, & Ding, 2019), which described the relationship between two sequences using a linear time-invariant system model. The syllable onsets were represented by a 0-1 sequence whose value was 1 only at the onset of a syllable. If the syllable onset sequence and the broadband envelope were referred to as *σ(t)* and *env(t)*, respectively. Their relationship was characterized using the following equation:

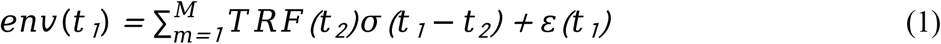

Where *TRF(t)* and *ε(t)* referred to the TRF and the residual error of the model, respectively. The order of the TRF model, i.e., *M*, was set to 200, corresponding to 1 s, and the pre-stimulus interval was set to 0.5 s. The TRF was estimated using ridge regression with a 10-fold cross-validation (Ding & Simon, 2012). The predictive power of the TRF was defined as the correlation coefficient between the predicted envelope and the actual broadband envelope, which was averaged over the 10 folds. When calculating the TRF of a speaker, all sentences from the speaker were concatenated.

### T5 modeling

A DNN model, i.e., Text-to-Text Transfer Transformer (T5, t5-base, Fig 6, (Raffel et al., 2020)), was used to estimate how well syllable onsets can explain the speech envelope when considering contextual information, i.e., the onsets of all syllables in a sentence. The syllable onset was characterized using a 0-1 sequence, the same as in the TRF analysis. The broadband envelope was normalized to integer numbers in the range of 0 to 100, where 0 and 100 are the minimum and maximum value, respectively. Both were down-sampled to 25 Hz with zero-padded to 500 points and then converted to tokens point by point with the tokenizer in T5 before fine-tuning by the “Trainer” from “transformers” (Wolf et al., 2019). The learning rate of the trainer was set to 1e-5, the training steps was set to 5000, and the training batch size was set to 32. The optimizer was adamw from “pytorch” (Paszke et al., 2019) with default hyperparameters, and a self-defined metric, i.e., predictive power, was used during fine-tuning to evaluate the training quality. Ten percent of the sentences were randomly chosen for validation, others were trained during fine-tuning.

**Fig 6.**
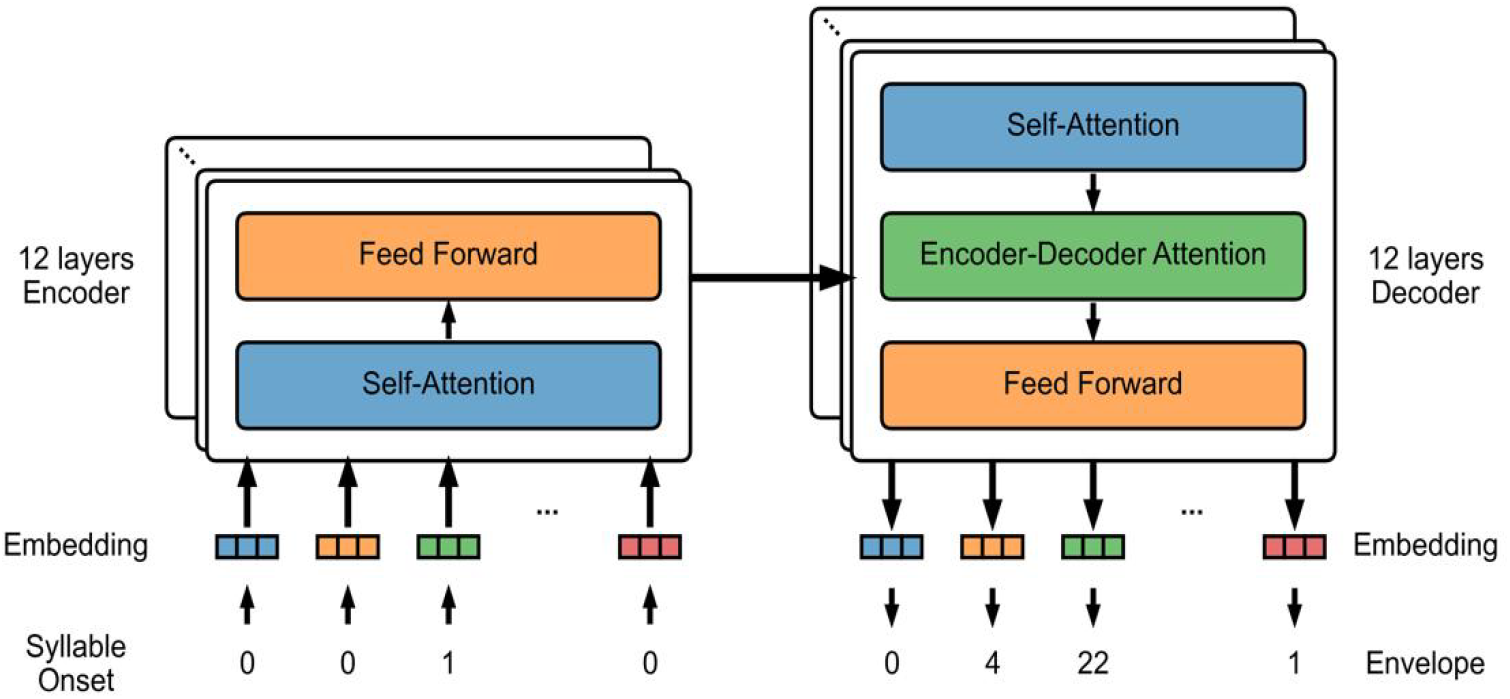
Architecture of T5 model used for sequence-to-sequence modeling.

## Supporting information

### MFA Validation

MFA Validation was carried out on TIMIT and Chinese-TIMIT, trying to test whether the syllable boundaries obtained by MFA were reliable. Syllable boundaries manually labeled by humans (referred as “manual” syllables) were compared to the syllable boundaries automatically extracted by MFA (referred as “MFA” syllables). The distribution of the rate of syllable duration in S1A Fig showed similar overall trends for both kinds of syllables, though there are slight differences. To judge this similarity, two statistical tests were further carried out. Firstly, the Pearson’s correlation, variance tests, and paired two-side T-tests were calculated between “manual” syllable rate and “MFA” syllable rate across speakers in S1B Fig. Results (for Chinese-TIMIT: r=1.00, p<0.01; v=0.00, p=0.97; t=0.66, p=0.51; for TIMIT: r=0.96, p<0.01; v=0.03, p=0.86; t=1.43, p=0.15) show high correlations with no significant difference.

**S1 Fig.**
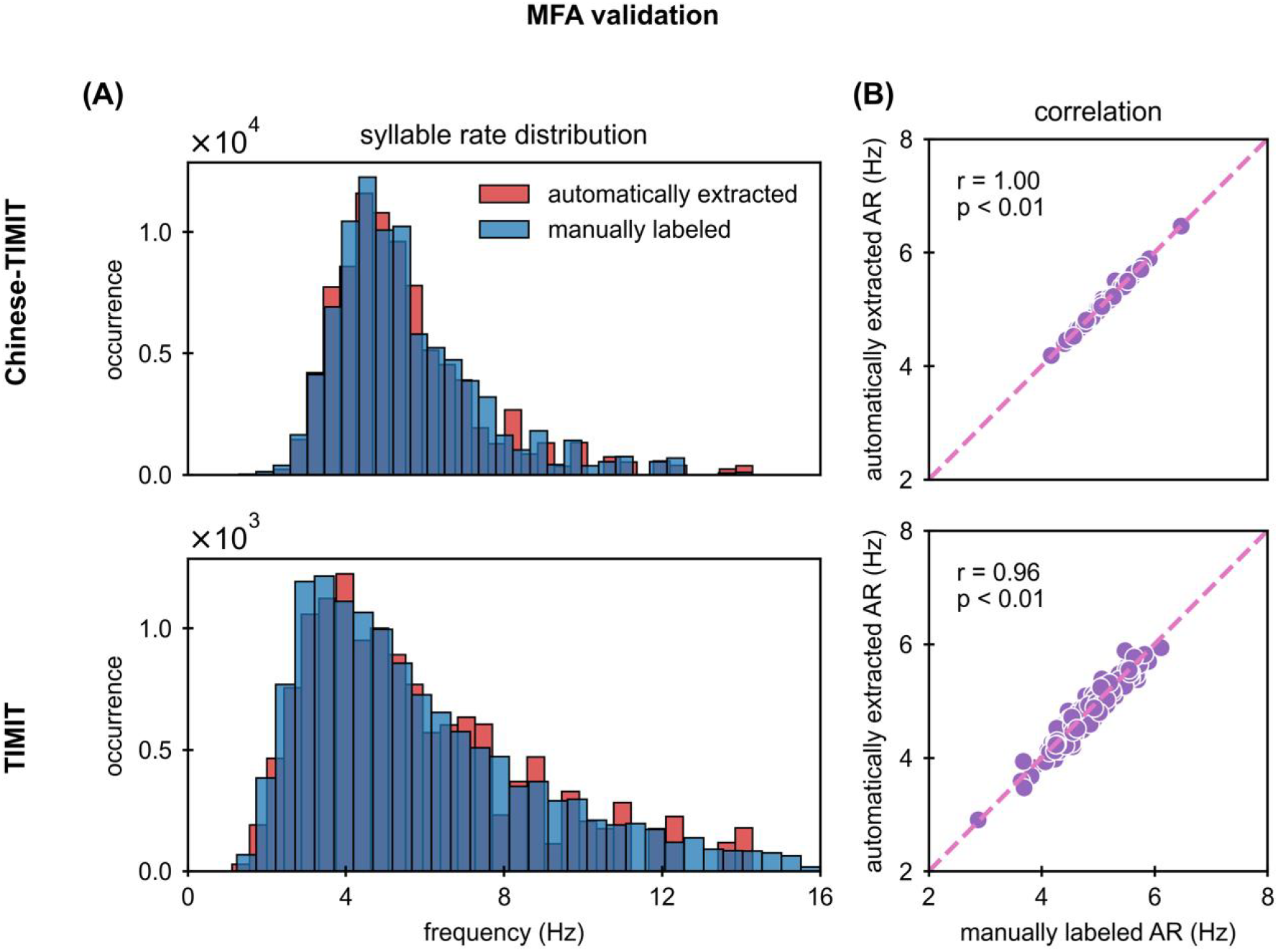
Validation of the syllable extraction method. (A) comparison of the syllable rate distribution based on the syllable boundaries that are manually labeled or automatically extracted by MFA. The two distributions are highly consistent. (B) scatter plot of the articulation rates of individual speakers that are calculated based on the automatically extracted syllable boundaries or the manually labeled syllable boundaries.

Having similar syllable rates does not mean having similar syllable onset sequences in the time domain. Thus, the other statistical test was based on TRF and bootstrap. The TRF was estimated by characterizing the relationship between speech envelope and different syllable onset sequences. In the bootstrap procedure, the “MFA” syllable boundary was shifted to left or right with a random time within 100ms to get “random” syllable boundaries. The TRF was estimated with a “random” syllable by 1000 times. The significance test is one-sided: if the “manual” syllable has higher TRF correlations with the “MFA” syllable than the “random” syllable in A times, the significance level is (A+1)/1001. As a result, the TRF correlations between “manual” and “MFA” syllables showed a high significance (P_TIMIT=0.011; P_Chinese-TIMIT=0.002), indicating that the “MFA” syllable onset sequences are very similar to “manual” syllable onset sequences and are reliable in the analysis.

### TRF for languages other than English and Chinese

The TRF analysis requires the extraction of syllable boundaries. For languages other than English and Chinese, the syllable boundaries were loosely extracted by grouping phonemes into syllables using the following rule: If a vowel was preceded by consonants, the onset of a syllable had one consonant. Otherwise, the syllable started with the vowel. For example, VCCCV was segmented into VCC and CV. Based on the syllable boundaries extracted this way, the TRFs were calculated for six languages and shown in S2 Fig. The results were consistent with the results in English and Chinese.

**S2 Fig.**
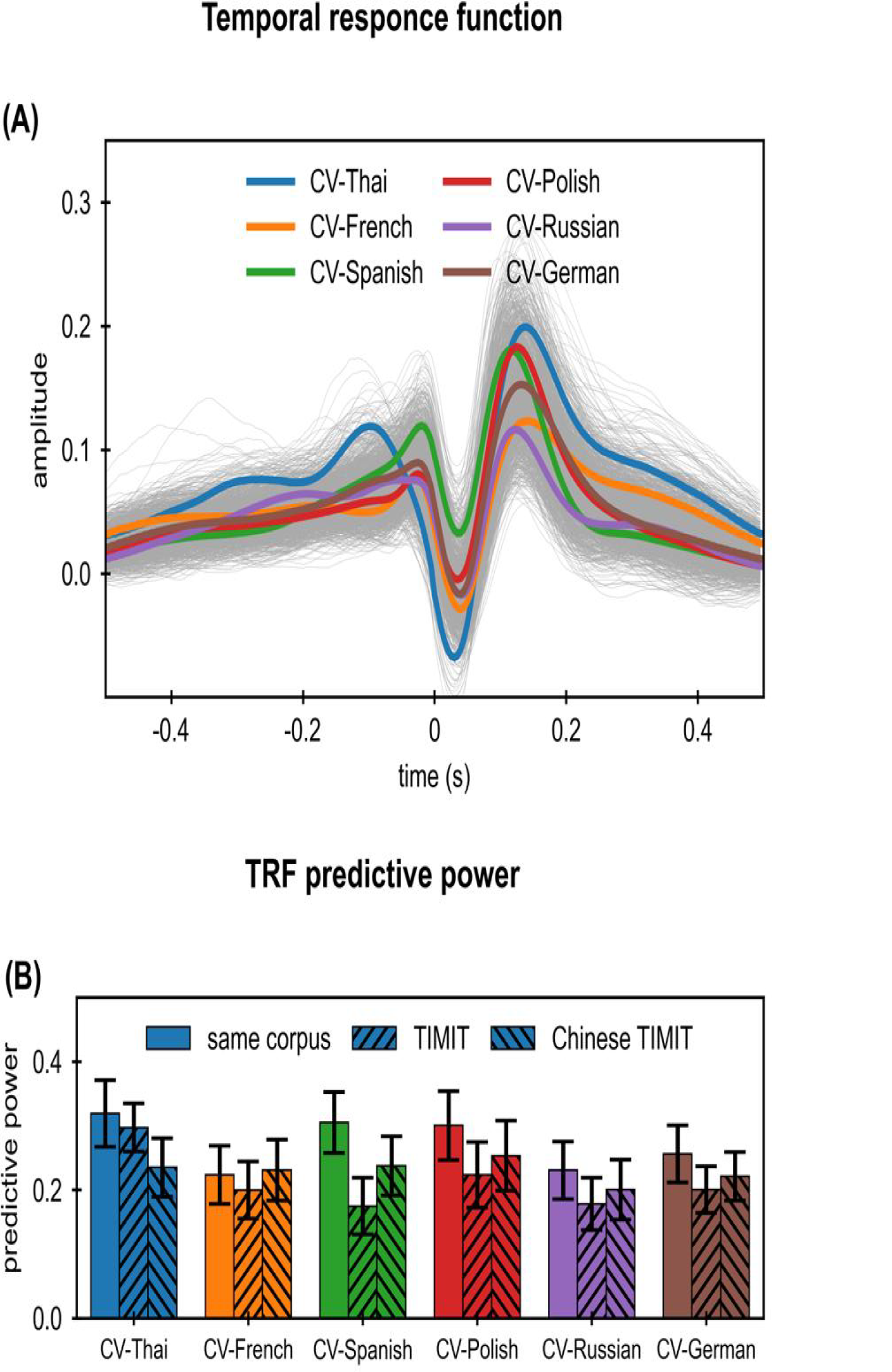
TRF for the Common-voice corpus. (A) TRF for each language is color-coded. The TRF for each speaker is shown by a gray curve. (B) The predictive power of the TRF for each language. The error bar shows one standard deviation.

**S3 Fig.**
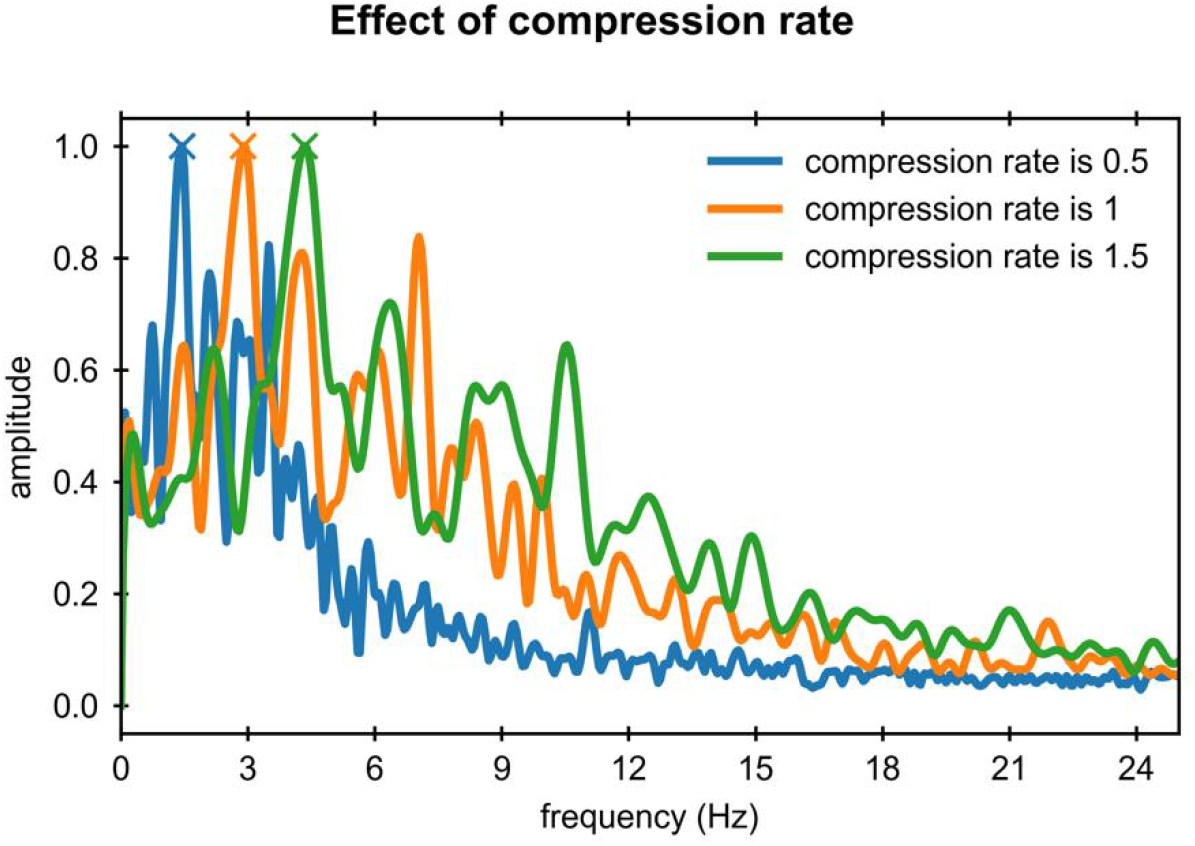
Modulation frequency shifts with the compression rate. The narrowband modulation spectrum was calculated with the sentence in Fig 1, and the peak modulation frequency was marked out by the marker.

**S4 Fig.**
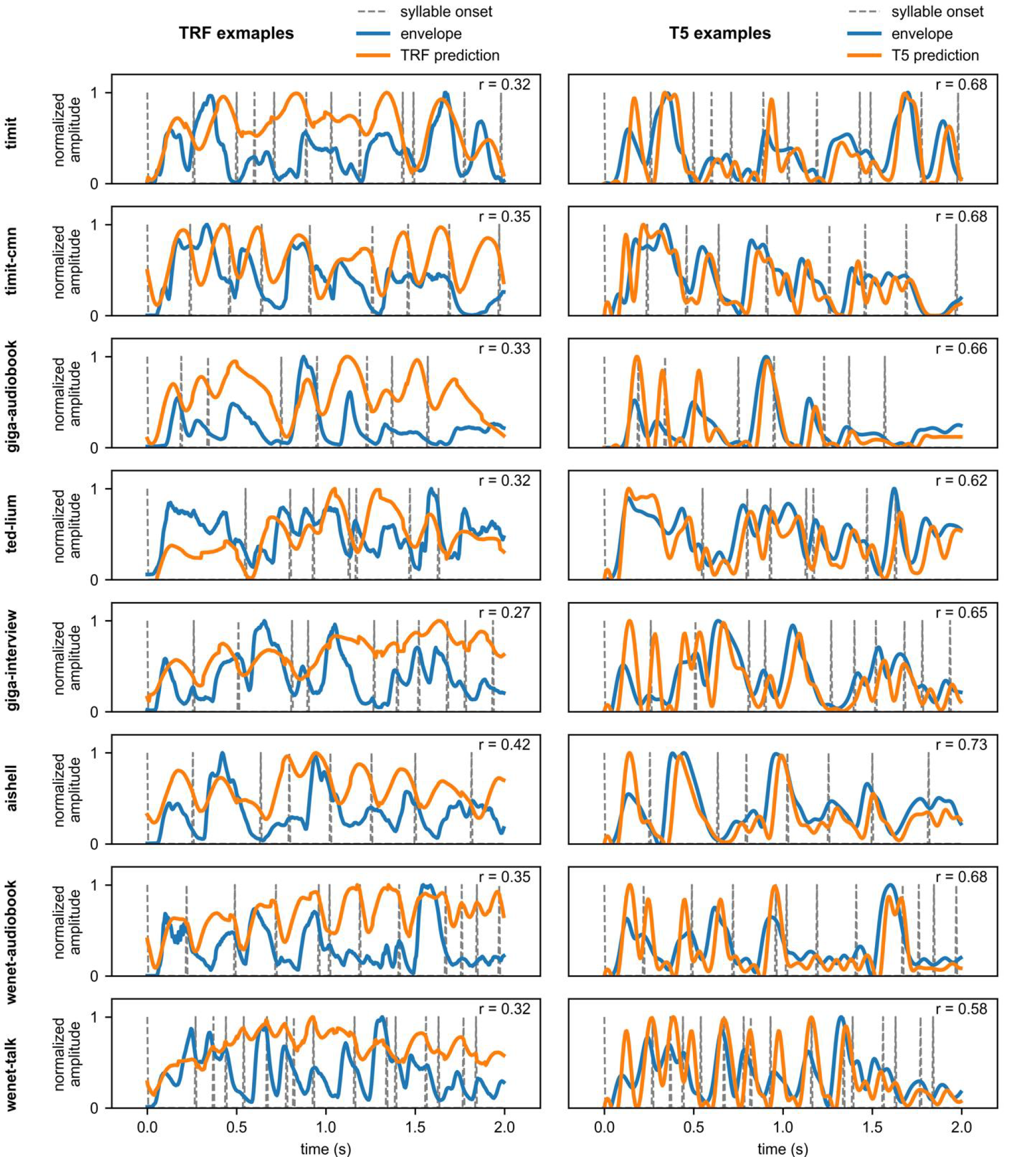
The examples of the predictions of the TRF and T5. Each line shows an example of a sentence from a corpus, which was cut off at 2 s. The sentences are randomly picked with the constraint that the predictive power of the sentence does not strongly deviate from the mean predictive power of the corpus for both models.

## Notes

### Competing Interest Statement

The authors have declared no competing interest.

